# MAARU: Multichannel Acoustic Autonomous Recording Unit for spatial ecosystem monitoring

**DOI:** 10.1101/2024.01.23.576628

**Authors:** Becky E Heath, Neel P Le Penru, James Skinner, C. David L. Orme, Sarab S Sethi, Robert M Ewers, Lorenzo Picinali

## Abstract

1.

1 – Acoustic localisation, which relies on simultaneous multi-microphone recording, adds spatial information to recorded audio and has been used in ecosystem monitoring to count individuals to improve abundance estimates, locate illegal activities such as logging/poaching, and monitor behaviour such as habitat use or species interactions. Studies have shown many advantages of acoustic localisation, but uptake remains limited as the equipment is often expensive, inaccessible, or only suitable for short-term deployments.

2 –Here, we present a low-cost, open-source, 6-channel recorder built entirely from commercially available components which can be integrated into a solar-powered, networked system. The MAARU (Multichannel Acoustic Autonomous Recording Unit) works in long-term autonomous, passive, and ad-hoc deployments. We introduce MAARU’s hardware and software and present the results of lab and field tests investigating the device’s durability, localisation accuracy, and other applications.

3 –MAARU provides multichannel data with similar costs and power demands to equivalent omnidirectional recorders. MAARU devices have been deployed in the UK and Brazil, where we have shown MAARUs can accurately localise pure tones up to 6kHz and 65dB bird calls as far as 8m away (±10° range, 100% and >60% of signals respectively), louder calls may have even further detection radii. We also show how beamforming can be used on MAARU devices to improve species ID confidence scores by 20%+ and recall by 10%+ when using the BirdNET automated bird identification algorithms.

4 –MAARU is an accessible, low-cost option for those looking to explore spatial soundscape ecology accurately and easily. Ultimately, the added directional element of the multichannel recording provided by MAARU allows for a new type of exploration into sonic environments.

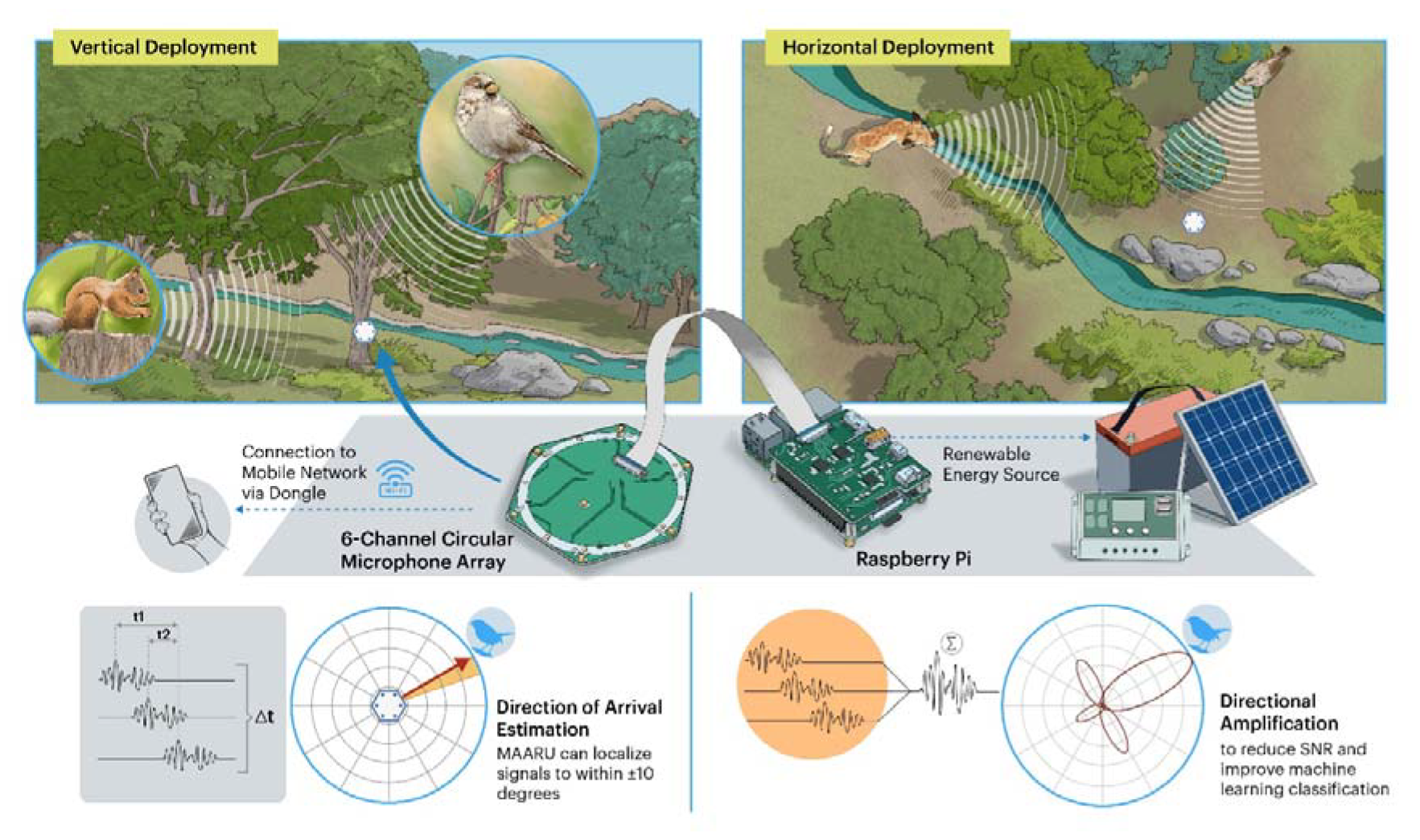

## 2. INTRODUCTION

Passive Acoustic Monitoring (PAM) is the use of Autonomous Recording Units (ARUs) to study ecosystem acoustics. PAM is cheaper and more scalable to large-scale, high-resolution wildlife surveys than human field data collectors (Darras et al., 2019), with reduced environmental impacts (Costello et al., 2016). Multi-microphone recording can add a new dimension of data from soundscape recording through acoustic localisation. Acoustic localisation uses slight differences in acoustic signals recorded from multiple points simultaneously to triangulate either a signal’s position (hyperbolic localisation) or direction of arrival (DOA) relative to the unit (Rhinehart et al., 2020).

Reviews of DOA estimation and acoustic localisation in an ecosystem monitoring context (Blumstein et al., 2011; Rhinehart et al., 2020) identify eight key purposes: assessing the positions and movements of individuals, studying interactions, identifying individual identities, sub-setting (and beamforming) sounds for more detailed analysis, calculating species abundance, and inferring habitat usage/territory. Additionally, acoustic localisation has been used to locate illegal logging and poaching (Andrei, 2015; Wijers et al., 2021). However, uptake of existing multichannel recorders remains limited as these devices are often non-or semi-autonomous, require specialist/discontinued equipment, lack full 360° localisation, or lack necessary hardware/ software for extended deployment in the field (Supplementary 1 -Bruggemann et al., 2021; Smith et al., 2021; Suzuki et al., 2017; Wijers set al., 2019).

Here we present an autonomous, self-powered and networkable recorder capable of DOA estimation and built from low-cost, off-the-shelf components. We demonstrate successful lab and field tests of MAARU and explain the software and hardware adaptations necessary for multichannel recording. Our device presents an accessible new option for autonomous multichannel acoustic recording.

## 3. DESCRIPTION

MAARU extends an existing omnidirectional (single-channel) Raspberry-Pi based, autonomous acoustic recorder (Sethi et al., 2018). We have modified this platform to incorporate multichannel recording for acoustic localisation by modifying: 1) the core recorder hardware, 2) onboard software, and 3) device weatherproofing.

### 2.1 Core Recorder Hardware

MAARU centres on a Raspberry Pi, a multichannel microphone array (Circular 6-Microphone Seeed Studio ReSpeaker array), a (renewable) power system and networking via a 3G/4G/LTE USB dongle (Figure 1). However, MAARU devices are also suitable for non-autonomous deployment with manual battery replacement and data download. The core MAARU set-up (Raspberry Pi and microphone array) weighs <1kg and measures 15×15×13cm with enclosure. The autonomous set-up weighs ∼12kg, with the largest component, the solar panel, measuring 84×58×15cm (Supplementary 3).

**Figure 1.**
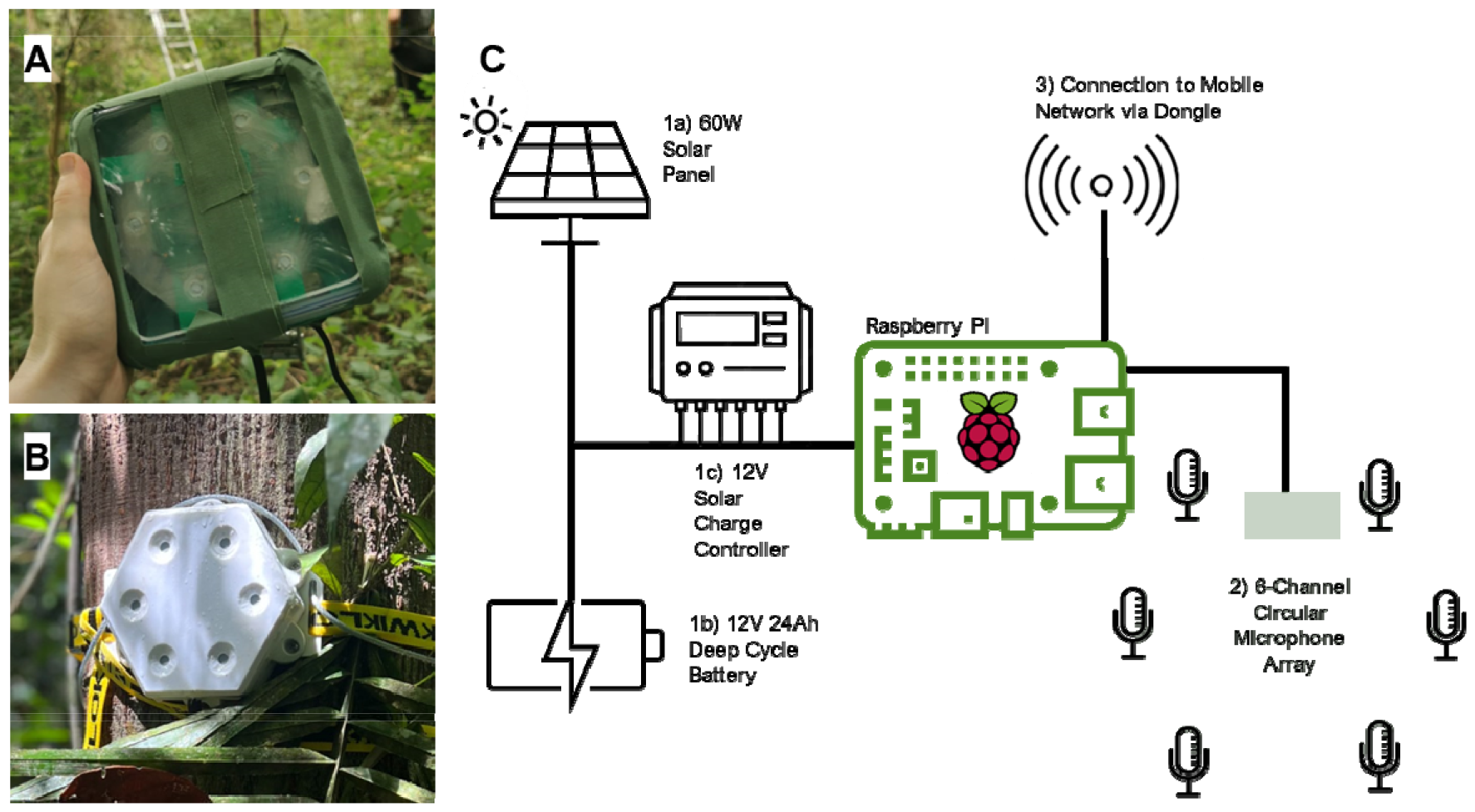
photographs of the MAARU recorder in the A) basic enclosure, and B) 3D Printed enclosure. C) Schematic of the MAARU uses a Raspberry Pi with the following attachments: 1) a renewable power source, (a) a 60W solar panel, b) a 12V 24Ah deep cycle battery, and c) a 12V solar charge controller. 2) A ReSpeaker 6—Mic circular array connected via the ReSpeaker multichannel interface. 3) a mobile dongle with a pre-paid mobile network sim card.

The ReSpeaker array uses six small (3.76 x 2.95 x 1.1mm), omnidirectional, Micro-Electro-Mechanical System (MEMS) microphones, fixed in a circular array on a PCB, which attaches to the Raspberry Pi via a Seeed Studio Multichannel HAT. We mediate clock drift (which is usually between 1 and 10 seconds a day Thode et al., 2006)) by managing all microphones with the same internal clock (sound card), ensuring the highly accurate and consistent synchronisation required for localisation (Mennill et al., 2012).

We found that MAARU uses 5W during simultaneous recording and upload, statistically not different to a single microphone recorder (Supplementary.4). A 5W device requires a minimum 120Wh per day. MAARU’s powering needs can be tailored to their environments (mains power, wind, non-recharging battery deployment). Powering equipment should have sufficient specification to generate more than the minimum allowance as given specifications are based on 100% efficiency which is not representative of real-world deployments.

One MAARU recorder generates 192kB of (16kHz, 16-bit, 6-channel) data per second (1536kb), reducing to approximately 96kB (768kb) after FLAC compression. For data upload to keep up with recording volume, the minimum required upload speed is therefore 0.768Mbps. Like the powering system, network connectivity can be tailored to the need and availability of connection. MAARU devices can be manual data download or autonomous data upload. For our autonomous deployments, we used a 3G/4G/LTE dongle. However, other connection types, such as Ethernet, and i-Fi Adaptors, could be used instead if appropriate with only small modifications to the setup required.

### 2.2 Device Software

Like the omnidirectional recorder that Sethi et al. (2018) developed, the MAARU software records, optionally compresses, and transmits acoustic data on a user-defined schedule while receiving updates and sending log information daily. For MAARU, we developed this code base by adding a multichannel sound card, rolling back package versions for compatibility, freezing the Linux kernel version, adding a kill switch, and switching to lossless compression.

The microphones are set to record 16-bit at 16kHz, and generate bulky WAV files, and while lossy compression is acceptable for audible range omnidirectional recording (Heath et al., 2021), DOA estimation requires a much finer-scale comparison between signals. We, therefore, use the open-source Free Lossless Audio Compression (FLAC) format to reduce file sizes for upload while preserving the original signal.

Full details of software amendments, the open-source code, and step-by-step set-up instructions are available at https://[Author name - removed for peer review].github.io/MAARU/.

### 2.2 Weatherproofing

In our deployments, we kept power storage components (the battery and the solar charge controller) separate from the microphone array and the Raspberry Pi to avoid interference. The power storage components were deployed in drybag. To weatherproof the microphone array and the Raspberry Pi we trialled two enclosure types: a modified 450ml plastic container, and a custom 3D printed enclosure (figure 1A and 1B). In both cases, the microphone holes are covered by ePTFE Acoustic Vents, which block water droplets without significantly affecting acoustic transmission (Supplementary 3b).

#### 2.2.1 Impact on Sound Quality

We used the MATLAB digital signal processing (DSP) toolbox (MATLAB, 2010) to compare recordings of a fixed-position 50-20,000Hz sinus-logarithmic sweep for multiple devices, with and without weatherproofing. Weatherproofing caused an average loss in the acoustic signal amplitude of <20dB (Figure 2 A, B).

**Figure 2.**
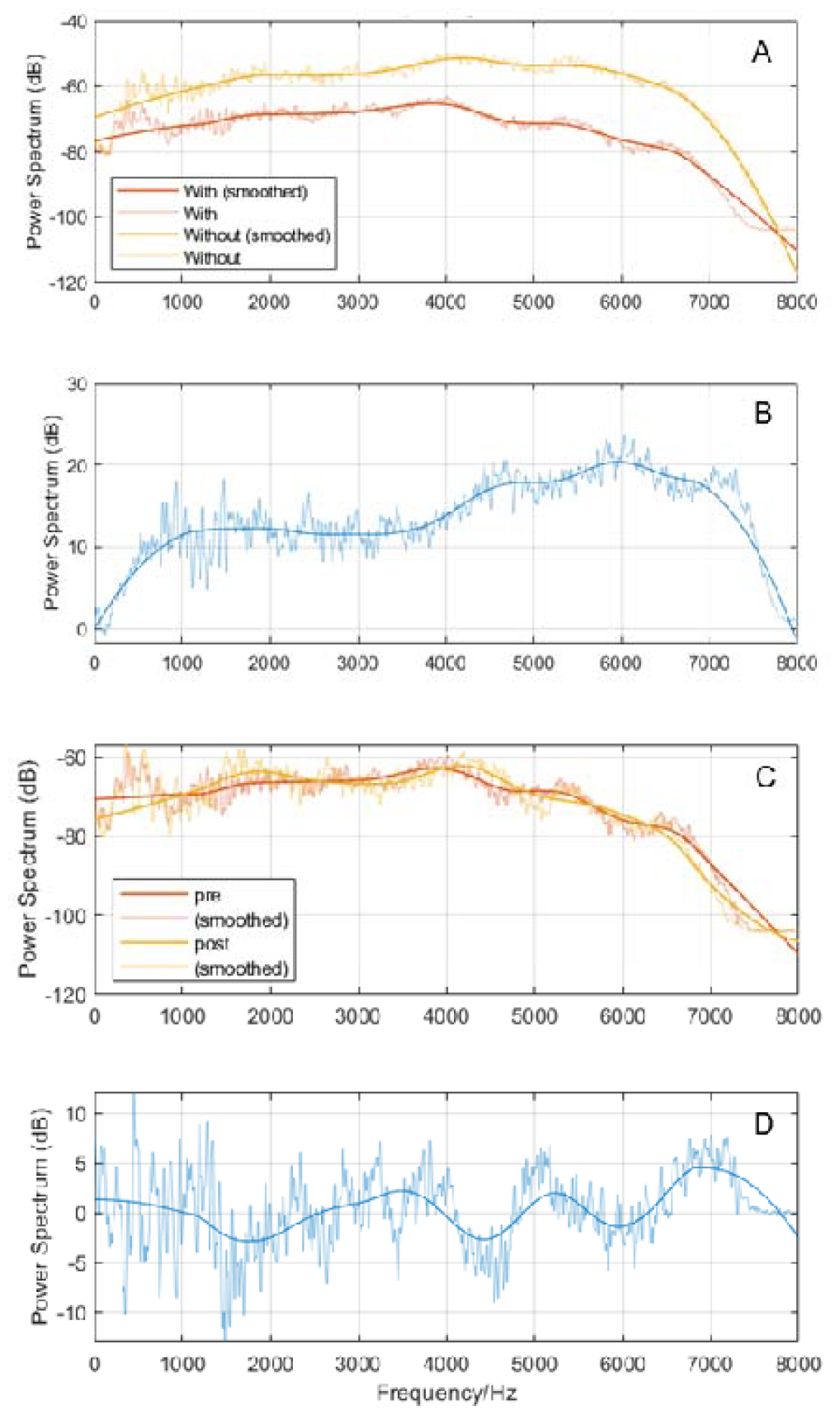
A) Power spectra of sinus-logarithmic sweep signals (sweeps) recorded from devices without weatherproofing (orange) and with (yellow line); B) the difference in the spectra in A; C) Sweeps recorded from devices pre- and post-6-month deployment in the field. D) the difference between the spectra in C.

#### 2.2.2 Durability

We have tested MAARU devices in four field trials/ experiments (Table 1). All devices remained watertight throughout deployments. Our longest deployment was a 6-months in Ancient Semi-Natural Woodland (ASNW) and Plantation on Ancient Woodland Site (PAWS) in the UK.

**Table 1.** outlines all the field deployments and experiments completed on MAARU recorders.

| Trial | Num. Recorders | Location | Duration | Type of Deployment | Deployment aim |
| --- | --- | --- | --- | --- | --- |
| Suburban Pre-Trial | 4 | London/ Kent, UK | 2 months | Fully Autonomous charging and upload | Test core functionality and basic weatherproofing |
| Temperate forest field trials | 4 | London/ Kent, UK | 6 months | Fully Autonomous charging and upload | Test long-term durability and basic weatherproofing |
| Temperate forest Field Experiments | 2 | Ascot, UK | 2 x 1 weeklong deployments | Non-Autonomous | Field test 3D printed weatherproofing |
| Rainforest Field Experiments. | 4 | Manicoré, Brazilian Amazon | Single day deployments, week and 1 month | Non-Autonomous | Test 3D printed weatherproofing<br>Explore beamforming |

In the Temperate Forest field trials, one recorder suffered an unrecoverable short circuit after a solar panel fell in a storm, and another failed after three months due to rodent damage on the power cable but ran as normal when re-powered. We have altered set-up instructions accordingly to recommend armoured cables and stronger straps. Data was retrieved successfully from all recorders in real-time until they failed. All recorders uploaded 24h data continuously in open-air pre-trials. However, partial panel coverage in the continuous UK trials led to data loss in autumn and winter, as daily sunlight could not compensate for power usage. We therefore advise further over-provisioning further for UK deployments. Despite this, 1700+ hours (550+ GB) of 6-channel audio were retrieved from the Temperate Forest field trials alone. Unfortunately, a kernel update mid-deployment caused a malfunction in the audio recording pipelines on-device, rendering some recordings unusable. We have since mitigated this problem by freezing the kernel. All four devices are once-again in working and currently being used in other experiments.

Post 6-month deployment, the devices were again tested with a 50-20,000Hz sinus-logarithmic sweep finding a ±5dB SPL difference in frequency response (Figure 2 C, D).

## 4. PROOF OF CONCEPT

### 3.1 Direction of Arrival (DOA) Estimation

Direction of Arrival (DOA) estimation is the prediction of the direction (azimuth) from which a sound originates relative to a device. Using a 2D array such as MAARU, this direction is represented by angles on a 2D plane between −180° and +180°, as illustrated in Figure 3A. The accuracy of MAARUs DOA estimation was tested in playback experiments, whereby test tones (pink noise, Eurasian Wren song (Karel, H. 2020), or single frequency tones) were played at 15-second intervals at five azimuths (−180°, −90°, −45°, 0°, and +90°) around a central MAARU device (figure 3A).

**Figure 3.**
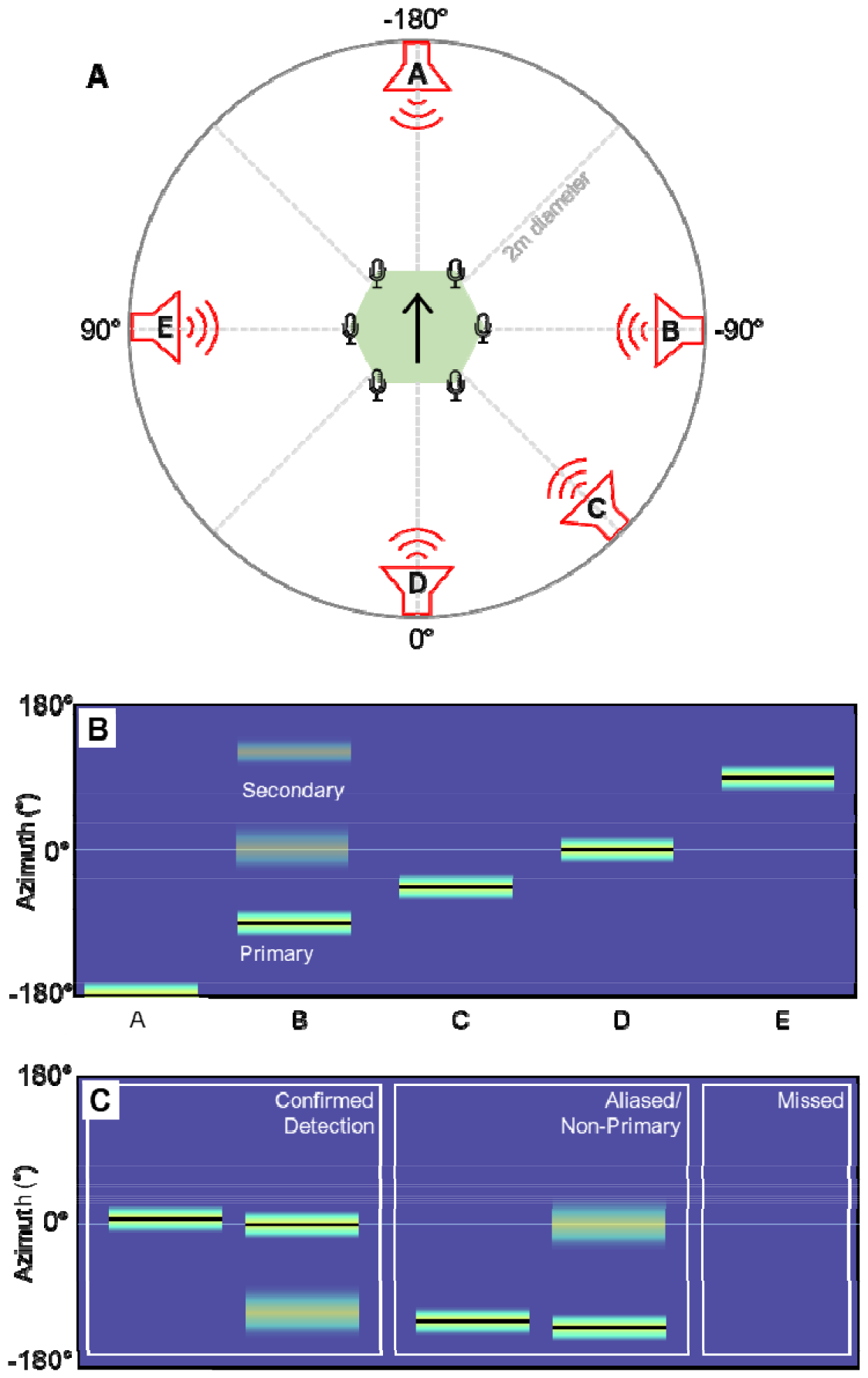
shows our DOA localisation test’s basic set-up and example outputs. A) an overhead schematic. B) an example direction likelihood matrix where all signals are localised correctly. For each time point, the lightness of pixels gives the MUSIC score (likelihood) that a sound is coming from a given direction (azimuth). The black line indicates the angles at which HARKBird determines there is a signal. Here, signals A-E are localised to the correct azimuth in order. Note: multiple positions can have high music scores and not count as a detection. C) illustrates the labels used to classify detections where the true azimuth is 0°.

We used HARKBird to estimate DOA (Suzuki et al., 2017), accessible at https://github.com/HARKBird-project/HARKBird. HARKBird adapts HARK software for robot audition: HARK (Honda Research Institute Japan Audition for Robots with Kyoto University) (Nakadai et al., 2017), which performs sound source localisation. HARK operates based on a transfer function (created using HARKTool5) and generates MUSIC (Multiple SIgnal Classification, Schmidt, 1979) power spectra for each time point. This spectrum is a one-dimensional vector quantifying the power of the MUSIC algorithm across all potential azimuths, which can be imagined as the likelihood a sound came from each possible azimuth. HARKBird is an intuitive user interface overlaying HARK with additional functionality for ecological applications.

In our experiments, we set the resolution of the transfer function to 5 degrees (i.e., 72 possible azimuths). In our configuration for each timepoint, HARKBird calculates the MUSIC power at all 72 possible directions, then, based on the given power thresholds, determines when a signal is present and the direction it has come from (Figures 3B and 3C). We then experimentally optimised MUSIC power threshold settings for each specific test (Supplementary 5) to account for irregularities in amplitude.

From HARKBird output figures, we determined whether a signal had been “detected”, “aliased/ non-primary”, or “missed” by inspecting the MUSIC matrix and determining the nature of detections (Figures 3B and 3C). Spatially aliased signals occur when microphones are too far apart, or sampling frequency is too low. Spatial aliasing can mean high-frequency signals return high MUSIC powers at multiple (inaccurate) points around the device, resulting in inaccurate DOA estimation.

The DOA estimation accuracy of MAARU was compared in four experiments (Table 2). Tests were conducted on weatherproofed (WP-Y) and un-weatherproofed (WP-N) devices at 1m unless otherwise stated. For all tests, we determined the azimuth error in °, proportion of signals with confirmed detections (figure 4).

**Table 2.** describes the localisation tests completed on MAARU as per protocol in Figure 4.

| Experiment | Groups | Origin | Test Location | Pre/ Post 6-month deployment |
| --- | --- | --- | --- | --- |
| Test Signal | Pink noise<br>Eurasian Wren | Generated in Audacity.<br>Xeno-Canto | Lab | Pre |
| Weatherproofing | Weatherproofed (WP- Y)<br>Un-weatherproofed (WP-N) | Na | Lab | Pre |
| Distance | 1m, 2m, 4m, 8m | Na | Field – Temperate forest | Post |
| Frequency | 100Hz, 400Hz, 500Hz,<br>1000Hz, 2000Hz, 4000Hz,<br>6000Hz, 7000Hz | Generated in Audacity. | Lab | Post |

**Figure 4.**
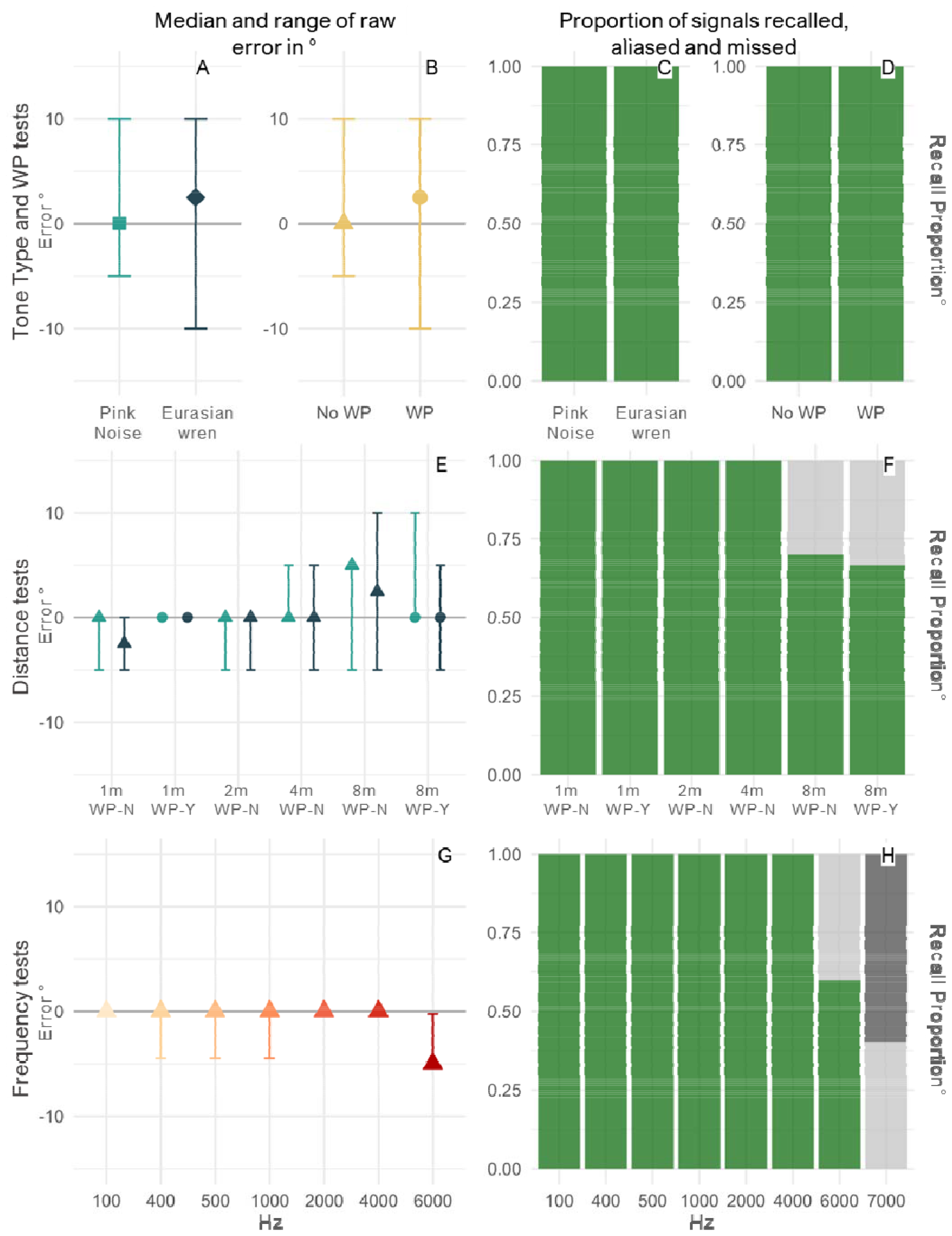
shows the outputs of all DOA localisation tests, as described in Figure 3. The left column shows the median and range of raw errors (differences between the True and Predicted azimuths). The right column shows recall/ detection rates (green, light grey and dark grey for detected, non-primary, and missed, respectively). The first row shows the results from lab tests at 1m, the second row shows the results from field distance tests, and the third row shows pure tone lab tests at 1m.

Across all tests, all *confirmed* detections were accurately localised to within ±10°, with the median error mostly being 0°, and not increasing to more than 5° (Figure 5). Recall remained at 100% for all signals except at 8m, 6kHz, and 7kHz (66.7-70%, 60% and 0% respectively). The only test in which no signals were detected accurately was the 7kHz single-frequency test (Figure 5 J). This loss of accurate detection may be due to spatial aliasing, however more investigation and modelling of the array is needed.

**Figure 5.**
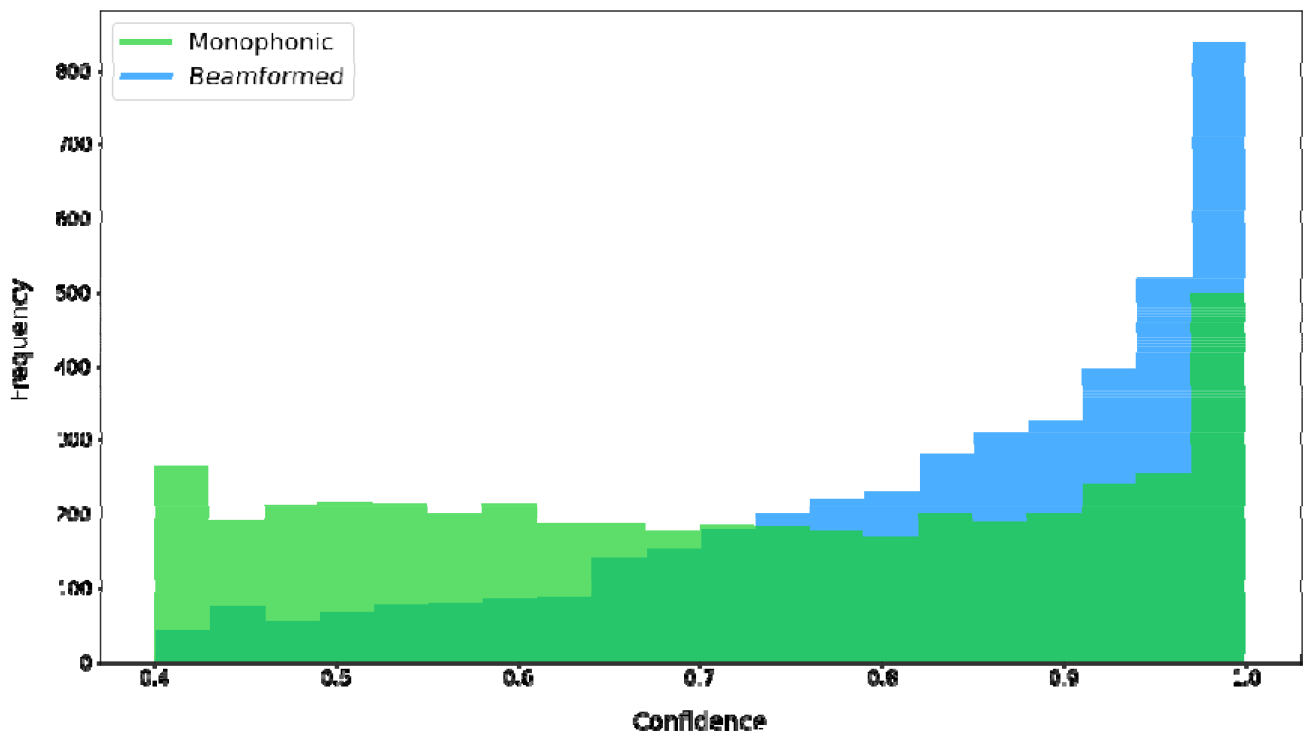
shows a histogram of confidence distributions across all detections for both monophonic and beamformed recordings. Beamformed recordings result in higher BirdNET confidence scores than monophonic recordings.

### 3.2 Beamforming for more accurate machine learning classification

Beamforming is the manipulation of time delays between recorded signals to amplify sounds that come from target directions. ODAS (Open embedded Audition System (Grondin et al., 2022)) is a robot audition platform, like HARK, which can be used for localisation and beamforming. Here, we used ODAS’ delay-and-sum beamformer to test whether audio beamformed to the known direction of origin could improve BirdNET classification over omnidirectional audio.

BirdNET is a deep neural network (DNN) that has been trained on 230,000+ labelled recordings (Kahl et al., 2021). The BirdNET takes in audio clips and returns bird species predictions with associated confidence values between 0 and 1. BirdNET is widely used within the bioacoustics community and has been used to classify over 140 million audio recordings (BirdNET, 2023).

#### 3.2.1 Improvement in confidence levels

We tested 55.1 hours of acoustic data from Manicoré (Brazillian Amazon) and 79 hours of data from Ascot (UK), both 16-bit, 16kHz. Beamformed signals were extracted for four directions relative to the recorder (0° azimuth, +45° Elevation; 0° azimuth, −45° Elevation; +45° Azimuth, 0° Elevation; −45° Azimuth, 0° Elevation).

For the confidence analysis, we only included species with that had been detected over 30 times in both monophonic and beamformed recordings (n=14). Descriptive analysis showed that the mean confidence using beamforming was higher than that of monophonic recordings (Figure 5, 0.83, SD= 0.15; 0.71, SD=0.19). The median confidence for beamforming was also higher than monophonic recordings (0.872 to 0.718 respectively). Wilcoxon signed rank test analysis showed that this difference was significant (W = 8457060.0, z = 44.64, p < 0.001).

#### 3.2.2 Improvements in Precision and Recall

We further tested whether beamforming could increase the precision and recall of BirdNET classifications using a 31-speaker array in a soundproofed lab (Le Penru, et al., 2023). In the test, 30 speakers played cricket noises at increasing amplitudes, whilst one would play Eurasian Blackcap or Carolina Wren. The amplitude of cricket noises was increased to create a gradient of Signal-to-Noise Ratio (SNR) from −15 (crickets louder than bird call) to +15 (bird call louder than crickets) in 5dB steps where both call types were as loud as each other at 0. The precision, recall and F_0.5_ of BirdNET classifications were calculated (Figure 6A, Supplementary 5).

**Figure 6.**
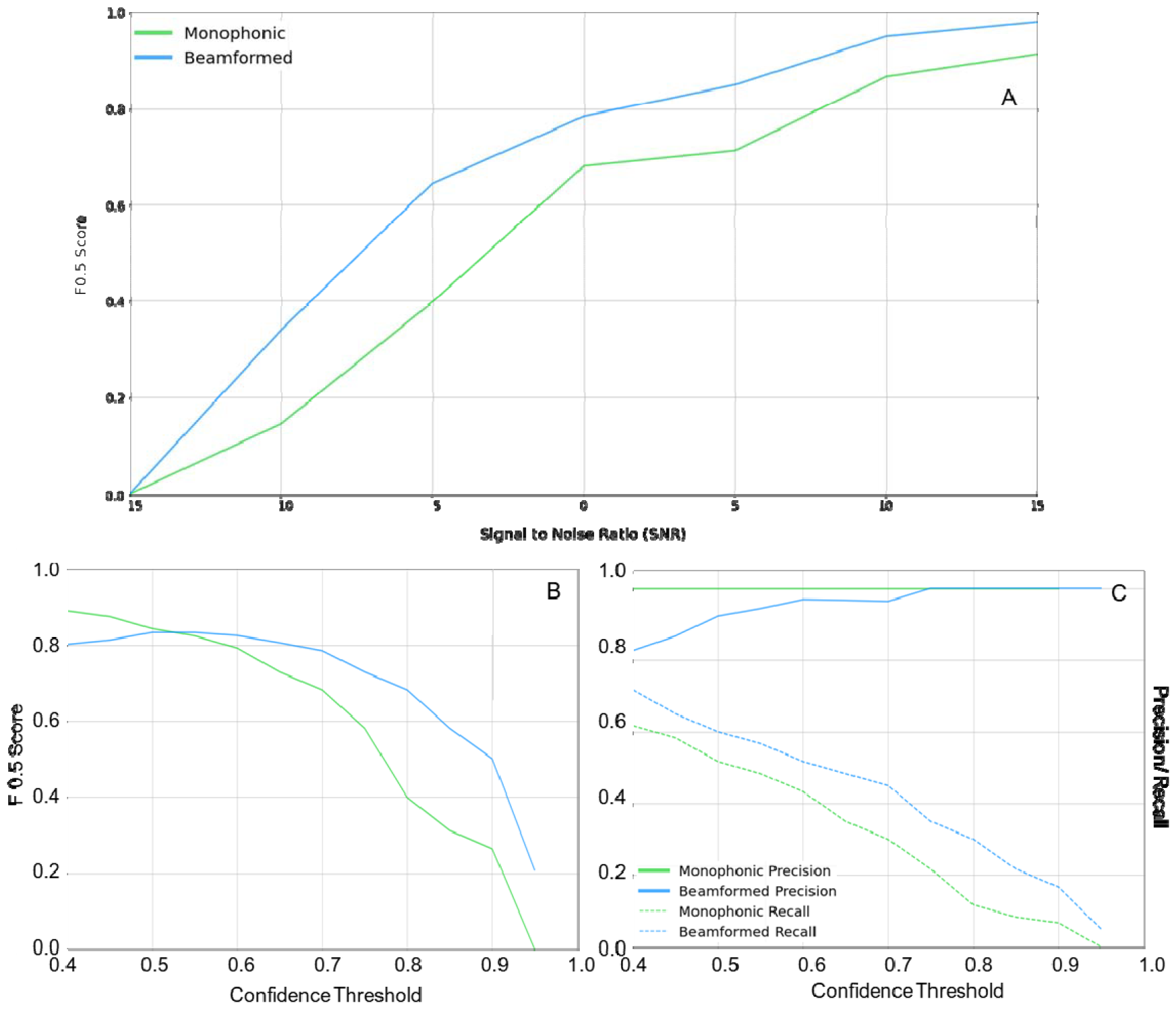
shows the response of BirdNET to beamformed and monophonic recordings. In A), beamformed recordings consistently achieve a higher F^0.5^ than monophonic recordings at a constant confidence interval of 0.7. In B), F^0.5^ for beamformed recordings surpasses monophonic recording beyond a confidence interval of 0.6. C) monophonic precision is consistently 1.0. However, by a confidence interval of 0.75, these values have equalised and beamformed recordings have consistently higher recall across all confidence thresholds.

At SNR=0, the beamformed recall was greater than monophonic recall (mean improvement of 0.11, SD=0.04), but precision was higher for monophonic recordings until a confidence threshold of 0.7, where subsequent thresholds yielded a precision of 1 for both (Figure 6C). Monophonic recordings yielded a better F_0.5_-score until a confidence threshold of 0.55, beyond which beamformed recordings provided a better score (Figure 6B). Overall, the mean improvement in F0.5-score after beamforming was 0.10, SD =0.12. Using a fixed confidence threshold of 0.7, beamformed recordings yielded a better F^0.5^-score at all SNRs, with the largest improvement at −5 dB (+0.25 score) (Figure 6A). Overall, the mean improvement in F^0.5^-score after beamforming was 0.12 (SD=0.08).

## 4. DISCUSSION

MAARU was designed to solve many commonly cited reasons for the lack of uptake of spatial acoustics in sound ecology (Table 3). MAARU combines state-of-the-art hardware and robot audition software to create a platform for enhanced ecosystem listening. MAARU recorders are capable of recording up to 48kHz, however localisation on this array works only for low-frequency audible-range signals (< 6kHz), so we recommend configuring HARKBird to localise signals <6kHz for 16kHz recording. MAARU and HARKBird are compatible up to 48kHz so it may be that this range increases with increased sample rates, although further testing is needed. Prospective users of MAARU should examine the general frequency ranges of the species they are hoping to monitor and determine whether *any part* of their call falls into the MAARU working range at different frequencies, which includes a broad reach of animal groups. High audible range and ultrasonic calling individuals will not be localised effectively with this recorder. Recall of a 65dB signal fell to 70% at 8m, however all animals call at different amplitudes so it would not be meaningful to advise a distance cutoff, however, we suggest using a high MUSIC threshold to avoid incorrectly localising weaker signals.

**Table 3.** describes the key aims and the degree to which they have been achieved with the MAARU recorder.

| Aim | Outcome |
| --- | --- |
| Low cost | MAARU combines autonomous PAM with usable multichannel recording for £206, only £26 more than its omnidirectional predecessor. Powering is an additional cost that can be tailored to the field area. |
| Accessible | All components could be sourced online at the time of writing. The basic MAARU can be built with a power drill and screwdriver. |
| Accurate | All signals could be localised to a high degree of precision ($\pm 10^\circ$)* |
| Durable | All devices remained watertight throughout deployments in temperate and tropical ecosystems. The early failure of the two devices has been addressed. |
| Customisable | MAARU is a modular recording unit which can be altered to suit powering and network capabilities. |
| Useful for ecologists | We have shown successful ports to ODAS, HARKBird and BirdNET. Audio from MAARUs can be used to accurately localise signals*<br>Beamformed MAARU audio increases BirdNET classification confidence and recall. |
| Low maintenance | Autonomous MAARU recorders can be left in the field indefinitely. The core MAARU devices have the same set-up and maintenance burden as popular omnidirectional devices. |

\*with signal components within loudness and frequency restrictions.

MAARU recorders have been demonstrated here for use in 2D localisation. However, networked MAARU recorders time-sync after every upload, making them equipped for inter-device/ array synchronisation. Theoretically, two or more MAARU recorders can be used to triangulate a location of origin and determine a calling position of an individual rather than a direction – however, this remains an open opportunity for future work.

## 5. CONCLUSION

For acoustic localisation to gain traction in ecosystem monitoring, equipment is needed that is widely available, inexpensive, self-synchronising and low maintenance (Rhinehart et al., 2020). Here we present a device that fulfils these conditions whilst also being suitable for indefinite autonomous deployment, with capacity for analysis in almost real-time. Localisation of signals in the field can describe patterns that single-channel acoustic ARUs alone cannot, we can track more precisely the movement, interaction, and relation to the habitat of individuals. These considerations provide a more complete understanding of individual responses to environmental stressors and could therefore facilitate more strategies for protecting at-risk organisms and environments.

## 6. DATA ACCESSIBILITY

[Removed for peer review]

