## Supplementary for "MAARU: Multichannel Acoustic Autonomous Recording Unit for spatial ecosystem monitoring"

**Contents:**

1- Emerging Spatial ARUs
2- Device Costs (incl. Sethi et. al.)
3- Device Technical Information
   a. General Information
   b. IPRO Vent Details
   c. Basic and 3D Printed Enclosure
4- Powering Assay
5- Full Localisation Experiment Details and MUSIC Power
6- BirdNET Classification Experiment Details
7- Device setup information
8- References

**1. Recent Spatial ARUs**

| **Name** | **Hyperbolic/ DOA** | **Mics? ARUs?** | **Base recording unit** | **Cost (min cost for study)** | **Data Collection** | **Mic**  **Synchro-nisation** | **Powering** | **Ref** |
| --- | --- | --- | --- | --- | --- | --- | --- | --- |
| Wildlife Acoustics | Hyperbolic | 2 mic (1 as failsafe)  3+ ARUs | SM2+ | £700  (£2,100) | Manual | GPS | Battery | (Mennill et al., 2012) |
| Audio-Moth Dev | Hyperbolic | 1 mic  3+ ARUs | Audio- Moth | £75  (£225) | Manual | GPS | Battery | (Hill et al., 2018) |
| SAFE Acoustics | Hyperbolic | 1 mic  3+ ARUs | R Pi | £200 | Cloud | NA (not yet done) | Self-charging | (Sethi et al., 2018) |
| VoxNet | DOA + Hyperbolic | 4 mics  3+ ARUs | Custom | ? | Cloud | Network | Battery | (Allen et al., 2008) |
| CARACAL | DOA + Hyperbolic | 4 mics  3+ ARUs | Custom | £150  (£450) | Manual | ? | Battery | (Wijers et al., 2021) |
| MAGO/ Dev-Audio | DOA + Hyperbolic | 7/8 mics  3+ ARUs | Laptop GUI | ? | Laptop required | Cable | Laptop required | (Sumitani et al., 2019) |
| Crunchant et al. 2022 | Hyperbolic | 1 mic  4 ARUs | R Pi | ? | Manual | GPS | Self- charging | (Crunchant et al., 2022) |
| WASN | Hyperbolic | W mic  5 ARUs | R Pi  (all commercial hardware) | ? | Cloud (through gateway Node) | Network | Battery | (Bruggemann et al., 2021) |
| **This one (MAARU)** | **DOA** | **6 mics**  **1ARU** | **R Pi** | **£226**  **(£226)** | **Cloud** | **Short Cable** | **Self-charging** | **This** |

**2. Device Costs (Accurate as of July 2021)**

Multi-Channel Device with ReSpeaker Mic Array:

| **Item** | **Cost GBP** |
| --- | --- |
| **Powering** | |
| Battery * | 119 |
| 12v to 5v step-down converter | 9.33 |
| 10m of AWG10 cable (10m) | 21.06 |
| Hooks and Slings (x4) | 14.00 |
| 10L Dri-Bag | 5.99 |
| **Electronics** | |
| Raspberry Pi 4 (4GB) | 54.99 |
| Respeaker 6 Mic Array (incl. soundcard) | 36.20 |
| Huawei Mobile Dongle | 38.98 |
| Memory Card** | 14.99 |
| USB-USB extension cable | 8.99 |
| Sistema Tupperware | 2.50 |
| IPRO Vents (x6) | 18.00 |
| Sniper Tape | 14.99 |
| **General** | |
| Electric Tape | 4.99 |
| Sealant | 7.95 |
| Polystyrene | 0.00 |
| Silica Gel (each) | 0.29 |
| Kwik-Lok Tie (each) | 3.15 |
| **TOTALS** | |
| Powering Total | 255.37 |
| Electronics Total | 189.64 |
| **Whole Device Total** | **461** |
| **Whole Device (Excluding Powering)** | **205.63** |

* Solar panel and battery requirements are location (sunlight) dependant so costs will vary depending on latitude of field site (London, GB used here).

** Memory card size also variable, 128GB used here.

Omni-Directional Device described by Sethi et. al. (EXCLUDES POWERING)

| **Item** | **Cost GBP** |
| --- | --- |
| **Electronics** | |
| Rasberry Pi B | 35.27 |
| Rode SmartLav+ Microphone | 47.00 |
| TRRS to TRS audio jack splitter | 6.59 |
| UASNW1 USB audio card | 8.99 |
| Memory Card 64GB*** | 11.99 |
| Huawei Dongle | 38.98 |
| 1m USB Extension Cable | 3.30 |
| USB to Micro USB Cable | 4.69 |
| Dri-Box | 10.99 |
| Dry Bag | 5.99 |
| **General** | |
| Cable Ties | 5.39 |
| Kwik-Lok (x2) | 6.30 |
| **TOTALS** | |
| **Device Total (Excluding Powering)** | **185.49** |

**3. Device Technical Information**

3.1 General Information

| Array Part Dimensions | 15x15x13cm |
| --- | --- |
| Array Part Weight | <1kg |
| Microphone Details | (MSM321A3729H9CP)  3.76 x 2.95 x 1.1mm  omnidirectional |
| Solar Panel Part Dimensions | 84x58x15cm (+ suspension equipment) |
| Solar Panel Part Weight | ~11kg |
| Microphone Part Attachment Height | 2/3m |
| Solar Panel Part Attachment Height | Canopy where possible, dependant on Sun exposure |
| Total Weight | ~ 12kg |
| Total Number Recorders | 4 |

3.2 IPRO Vent Details

(Taken and adapted from [www.ipromembrane.com](http://www.ipromembrane.com))

Vents used in this device were the B0.2 as their properties most closely matched our requirements.

| IP Rating | IP67, IP68 (5m water immersion for 1h) |
| --- | --- |
| WEP (60s) | 300kPA |
| Maximum Transmission Loss  (Max Value 100 – 10,000 Hz) | <1.5dB |
| Material Type and Colour | White ePTFE |
| Characteristic | Hydrophobic |
| Reference Thickness (mm) | 0.4 |
| Temperature Range (^O^C) | -40 to 85 |
| Adhesive Type | Acrylic |
| Recommended Orientation | Internal Mount |
| ROHS Compliance | Yes |

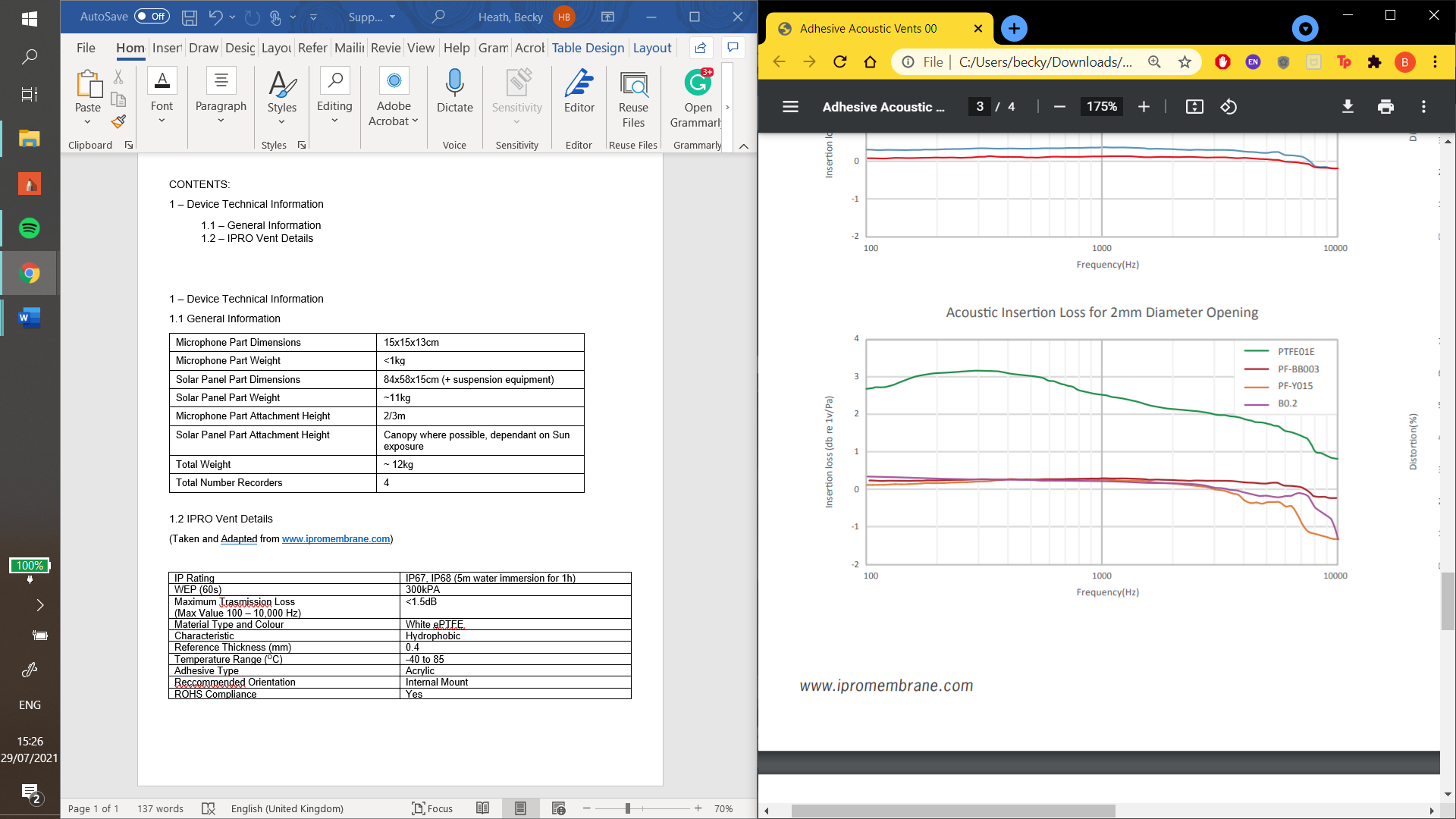

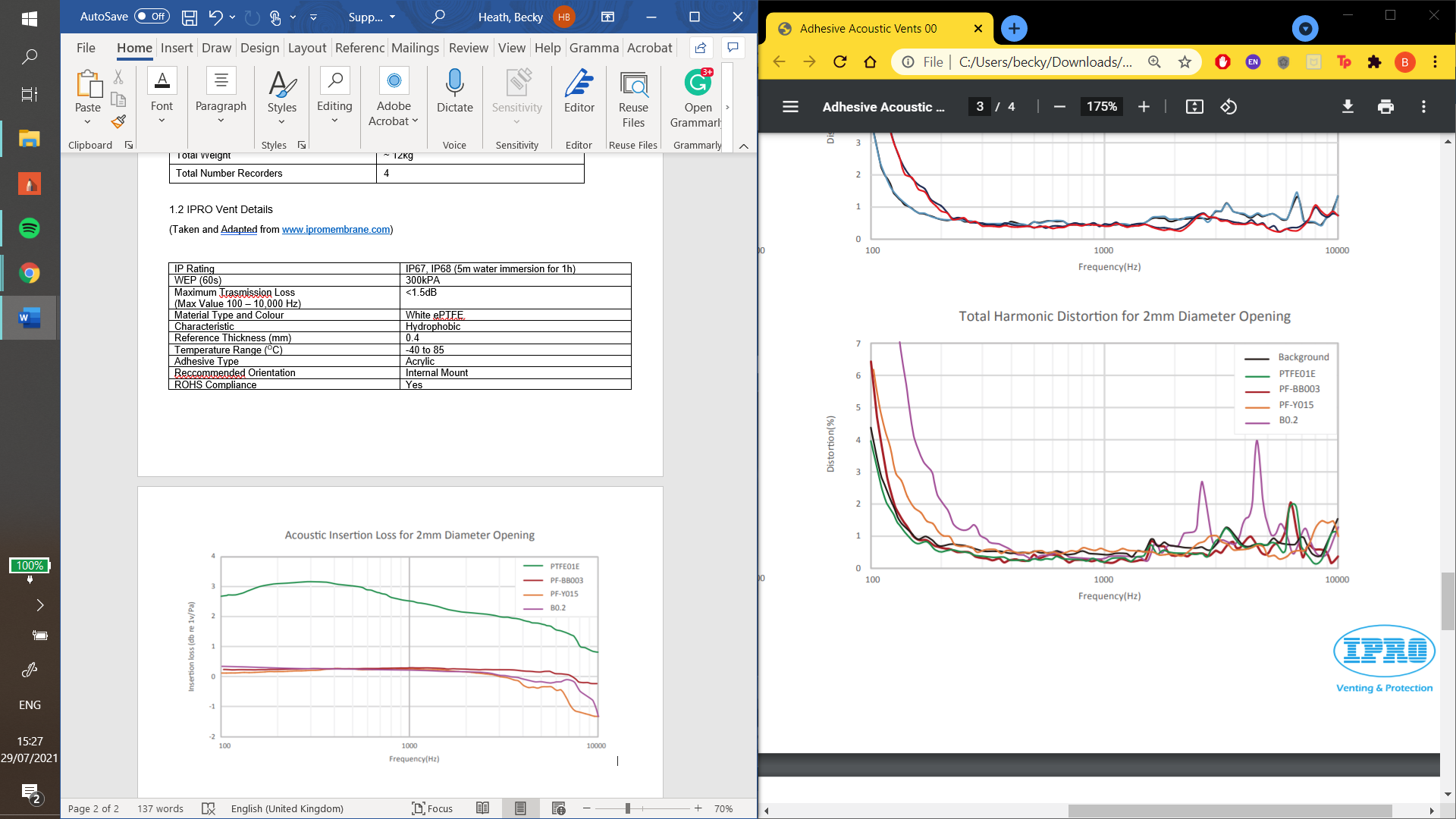

3.3 Weatherproofing Pictures

**Basic weatherproofing:**

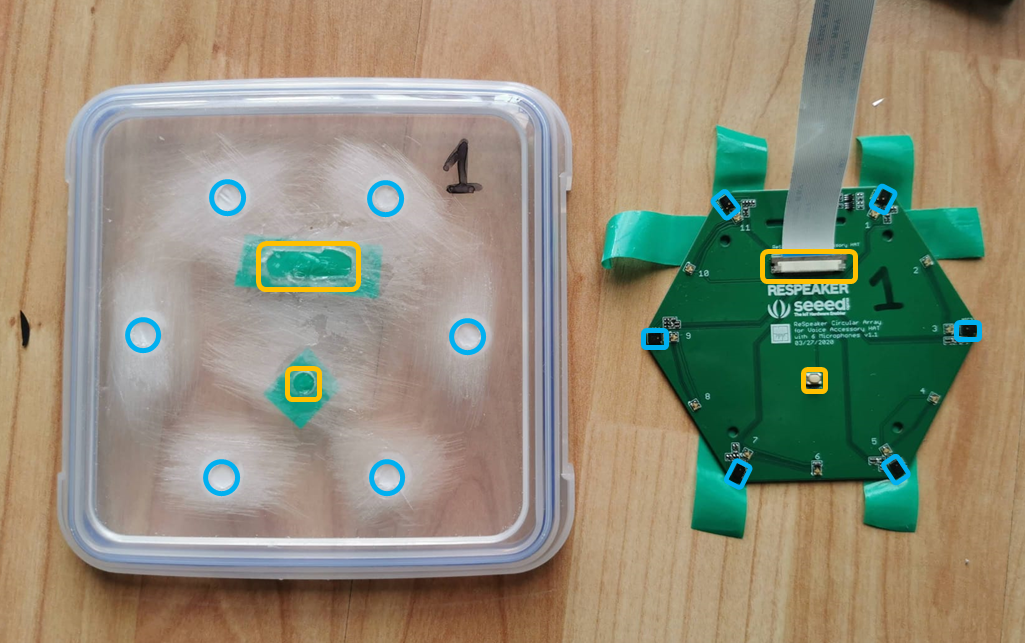

The sound-permitting device weatherproofing. The full blue circles correspond to holes in the container lid covered by ePTFE waterproof acoustic vents, and these holes line up with MEMS microphones (blue rectangles) on the array. The array is held flush to the container lid by additional holes made for other raised components on the PCB (yellow rectangles), which are instead sealed flexibly with an air-and-water tight sealant.

**3D printed weatherproofing**

Below shows the development process of the 3D printed housing for the MAARU recorder

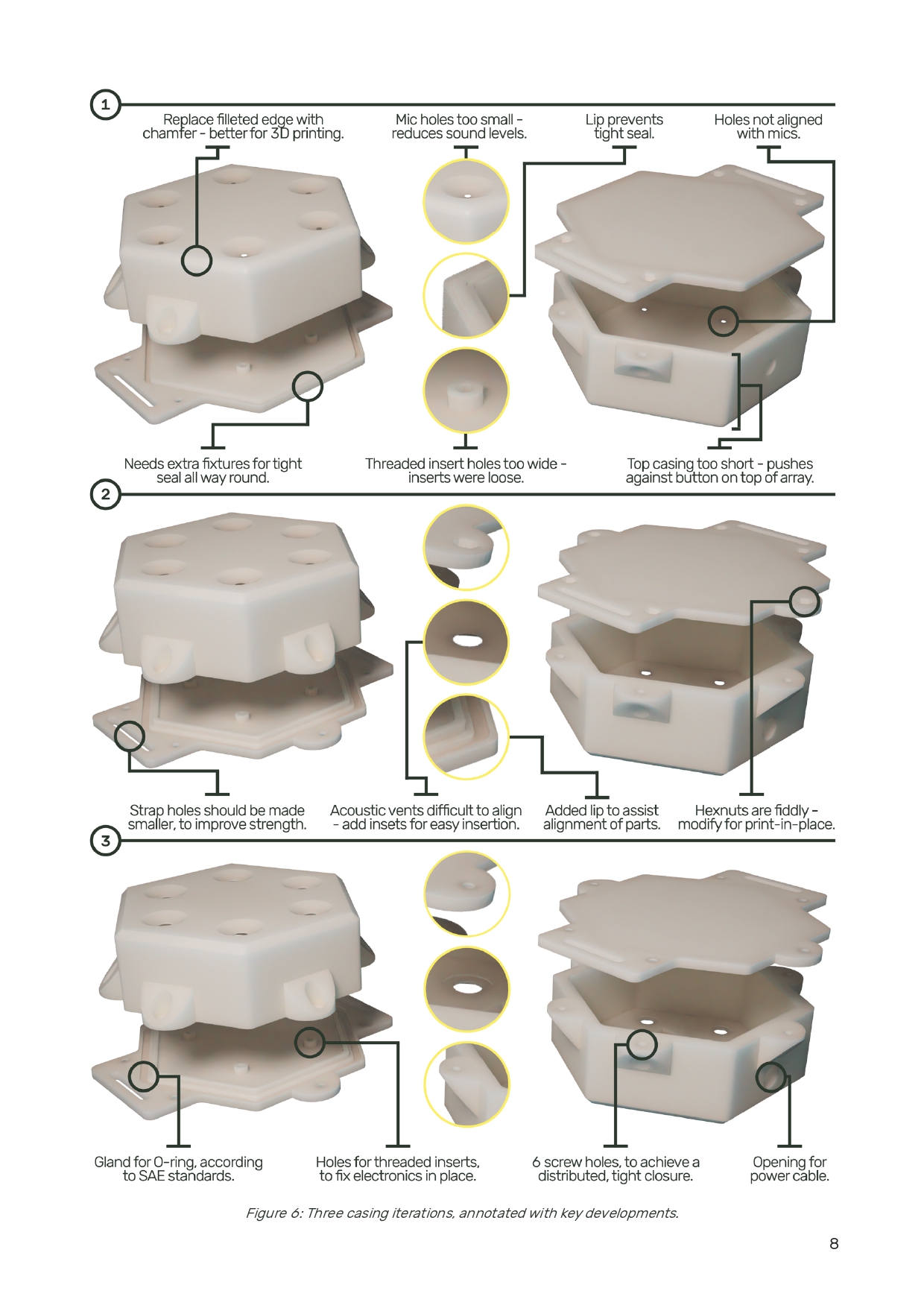

**4. Powering Assay**

Power Consumption

To determine the power usage of the multi-channel and omnidirectional devices, regular watt readings were taken from both devices (via a USB power reader) during continuous and non-continuous recording. In one test, a screen (showing status information) was attached to the recorder so the exact status of the device could be determined; in another, the device was left as it would be *in situ,* and an average reading was found. In both cases, readings were taken every 15 seconds for 5 minutes. We found that the addition of multiple microphones had no significant impact on the power usage compared to the omnidirectional device, in all phases: start-up (PO), recording (REC), and simultaneous recording and upload (REC+UP) (Respectively: t= 0.87_10.51_, p= 0.400, t= 1.88_20.5_, p= 0.075 and t= 0.253_9.39_, p= 0.8054, Welch’s T-Test.). Data transmission is almost instantaneous in the omnidirectional recorder, so no measured time points fell during a period where recording and upload were happening simultaneously in the non-continuous test.

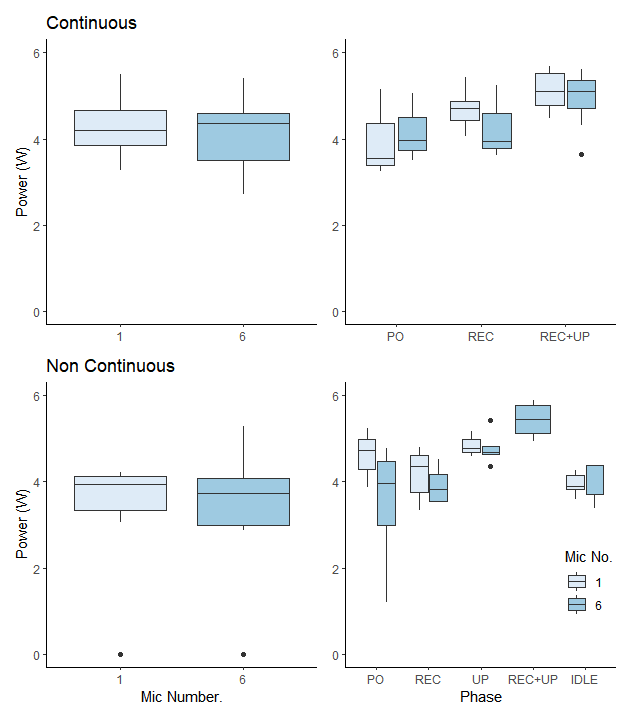

Shows the comparative power usage between the 1- and 6- Mic recorders under different running conditions. The left panels show values averaged over a 5-minute period. The right panels show the individual running periods separately PO = Powering on, REC = Recording audio data, UP= Uploading data, REC+UP = Simultaneous recording and upload. Power consumption appears slightly higher for both in the continuous recording. Black dots show outliers. Overall, there is no significant difference in power consumption between the devices (T >0.05 in all cases).

**5. Full Localisation Experiment Details and MUSIC Power**

| **Experiment** | **Test Group** | **MUSIC power threshold** | **Number of Tests** |
| --- | --- | --- | --- |
| Test Signal | Pink noise | 24 | 24 |
|  | Eurasian wren | 24 | 12 |
| Weatherproofing  (Basic) | Yes | 24 | 12 |
|  | No | 24 | 24 |
| Distance | 1m | 26.5 | 37 |
|  | 2m | 26.5 | 20 |
|  | 4m | 26.5 | 20 |
|  | 8m | 25 | 38 |
| Frequency | 100Hz | 27 | 5 |
|  | 400Hz | 27 | 5 |
|  | 500Hz | 27 | 5 |
|  | 1000Hz | 27 | 5 |
|  | 2000Hz | 27 | 5 |
|  | 4000Hz | 27 | 5 |
|  | 6000Hz | 27 | 5 |
|  | 7000Hz | 27 | 5 |

**6. BirdNET Classification Experiment Details**

Below is the precision, recall and F_0.5_ Score calculation equations:

$$Precision= \frac{TP}{(TP+FP)}$$

(1)

$$Recall= \frac{TP}{(TP+FN)}$$

(2)

$$F_{0.5}= \frac{Precision \times Recall}{\left( {0.5}^{2} \times Precision \right)+Recall}$$

(3)

**7. Device Setup Information**

From the downloadable resources from https://beckyheath.github.io/MAARU

**Hardware basics:**

| 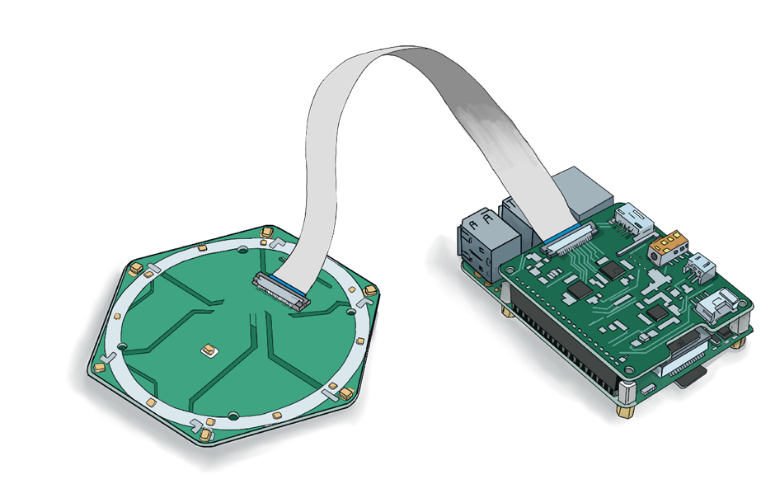 | |
| --- | --- |
| 1. Connect the ReSpeaker 6-microphone array to the ReSpeaker HAT via the ribbon cable as above. | 2. Attach the HAT and the array to the Pi via the 40-pin header on your raspberry Pi. |

**Setting up the operating system:**

There are two options for setting up the operating system. 1) you can download our prebuilt OS [[here]](https://drive.google.com/file/d/1sTKPgUOcT4SQeJqtF6wdd6rjtqFLZzfR/view) on to a micro-SD card or 2) follow instructions to set it up yourself [[here].](https://github.com/BeckyHeath/multi-channel-rpi-eco-monitoring)

| 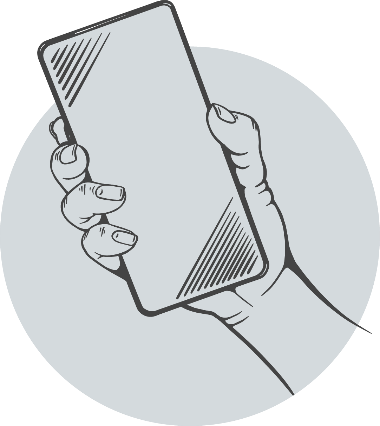 | |
| --- | --- |
| 3. You’ll need Wi-Fi or a network connection to get everything set up. If you’re looking to autonomise your device by connecting it to mobile networks, you can set that up now: | |
|  | **For networked recorders….**  a. Pay for a network subscription and activate your SIM card. SIM card activation usually involves making a phone call from your phone with the SIM inserted but can vary network-to-network. Make sure you can connect to the internet via the SIM on your phone first.  b. Put the SIM in the USB dongle.  c. Plug the USB dongle into the Raspberry Pi |
|  | **For offline recorders…**  You’ll want to use WiFi or ethernet to get the Raspberry Pi |

**Once you’ve got a functional Operating system on a micro-SD:**

| 4. Insert the micro-SD to the Raspberry PI | 5. Plug the Raspberry Pi into a screen and power on. Check that startup works as expected | 6. Power down******. Unplug the screen (and keyboard and mouse if using) and you’re good to go!! |
| --- | --- | --- |

**TIP**: Be careful not to suddenly unplug the raspberry Pi at any point. It’ll mess with the system. Instead, if you want to turn off the Pi:

- Press the button in the center of the array and wait for it to power down.
- Type “sudo halt” into the command line.
- Switch off the power source (wall plug or solar charge controller).

**If you want to power MAARU autonomously...** check the autonomous powering document.

**If you don't want to power MAARU autonomously...** you should plug your pre-charged battery into a solar charge controller. Or if you are chosing to use a power bank with inbuilt charge moderation, you can plug right into the Pi

**Autonomous Powering**

Here we explain how to power your MAARU autonomously with solar power. If you have mains connection available, you can just plug into the wall. Similarly solar panels can be subbed for wind or other renewable power sources.

**If you don't want to power MAARU autonomously...** you should plug your pre-charged battery into a solar charge controller. Or if you are chosing to use a power bank with inbuilt charge moderation, you can plug right into the Pi

**For solar powered deployments...**

|  | 1. Connect the AWG10 cable to the solar panel. You can do this either by stripping and soldering this to the existing wire or by attaching the new cables directly to the panel (Keep note of the negative and positive terminals). |
| --- | --- |
| 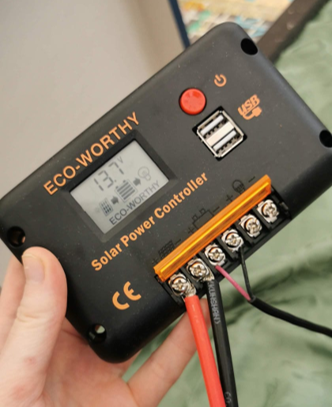 | 2. Connect the solar panel to the solar charge controller by the AWG10 wire. Ensure you connect positive to positive and negative to negative. Left two terminals in the picture here. |
| 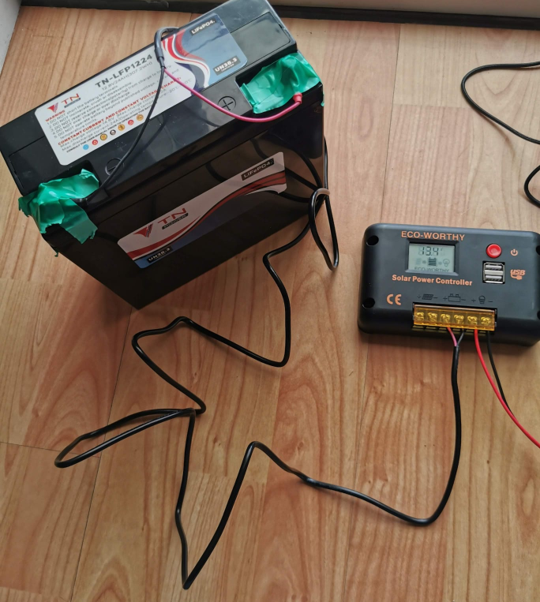 | 3. Connect the Battery to the solar charge controller (middle two terminals). You can use the AWG10 for this or I’ve often just used the wires that came with the panel/ solar charge controller set. Again, ensure you connect positive to positive and negative to negative. Note: the solar panel isn’t attached here, but if for your field deployments make sure that it is. |
|  | 4. Go through the settings on the Solar Charge Controller and make sure it all matches up to your battery. Especially: running voltage and battery type. This will maintain battery quality for longer |
| 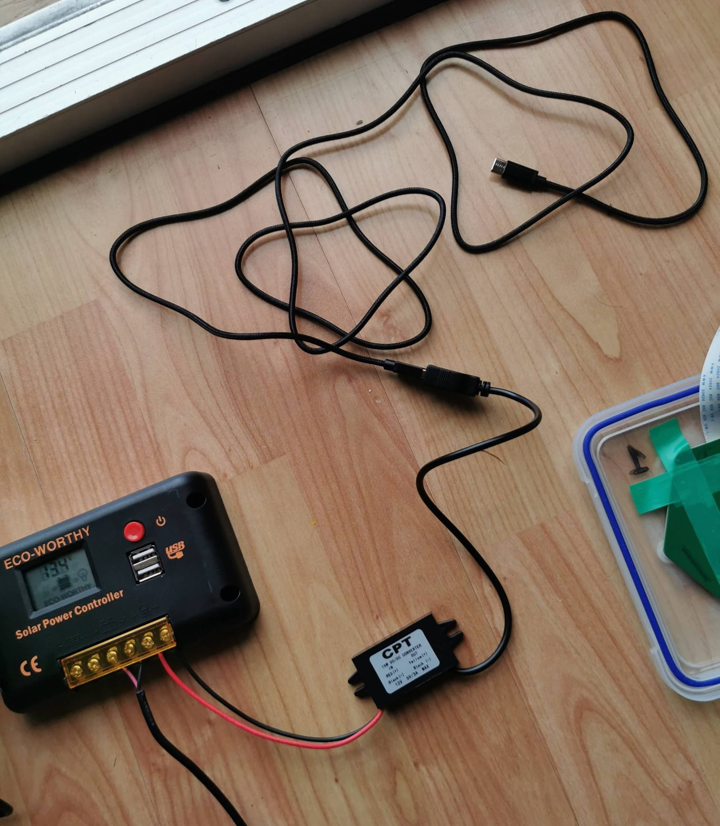 | 5. Attach the 12V to 5V step-down to the power out terminus on the Solar Charge Controller. Ensure the terminals match up and that the 12V side is attached to the solar charge controller. Right two terminals here. Note: Depending on the type of step-down converter you have you may need a USBC adaptor. |
| 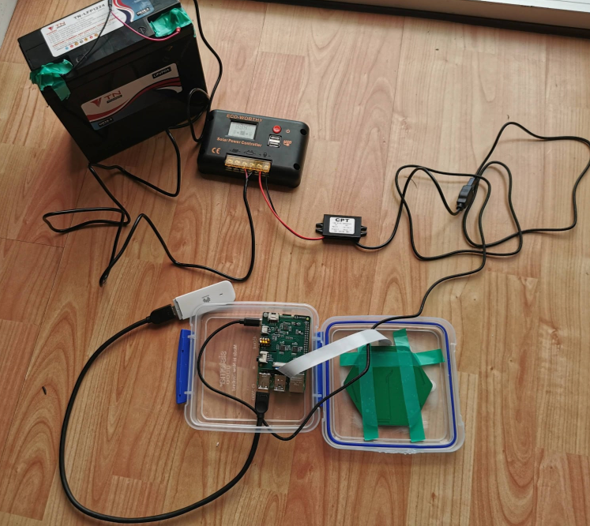 | 6. Plug the power source into the Pi |

**TIP:** You can control whether the power is running to the Pi or not by the button on the solar charge controller

**Basic Weatherproofing**

**Raspberry Pi and microphone array waterproofing.**

We present here two different methods of weatherproofing for MAARU, depending on whether or not you have access to a 3D Printer. For both, you’ll need acoustic weatherproof vents (details in kit list).

Tip: Power down and unplug the device from the power source while you’re setting up the waterproofing

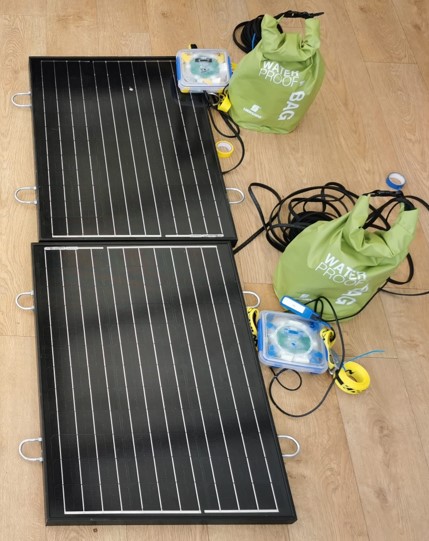

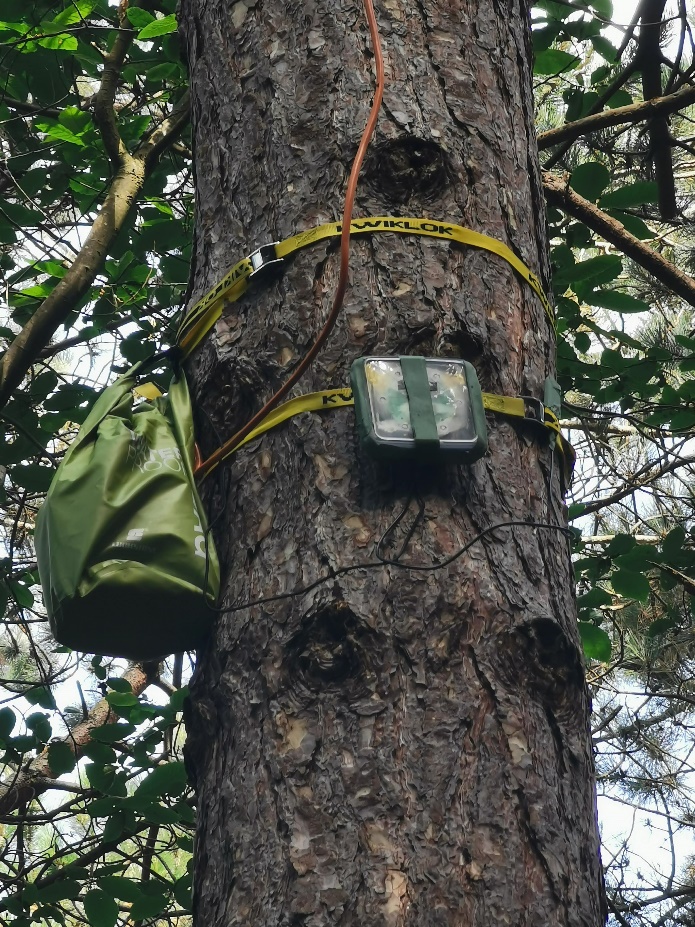

| 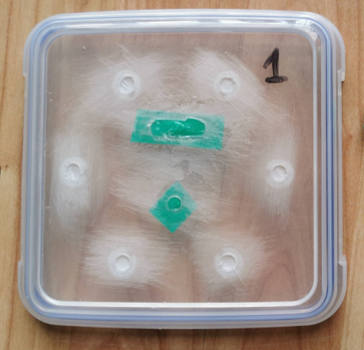 | 1. Drill a 1x3cm hole in the bottom of the box and 6+2 holes in the lid. The hole at the bottom of the main box will allow for power/data wires to exit the Tupperware. The 6 circular holes on top will line up with the MEMS microphones on the array while the 2 other holes will line up with two other protrusions on the surface of the array. Once all the holes have been made, sand down the edges and surrounding area to reduce risk of damage and increase surface area for adhesives to attach. Attach the IPRO vents to the mic hole and lay electric tape over the other two holes and cover with sealant. |
| --- | --- |
| 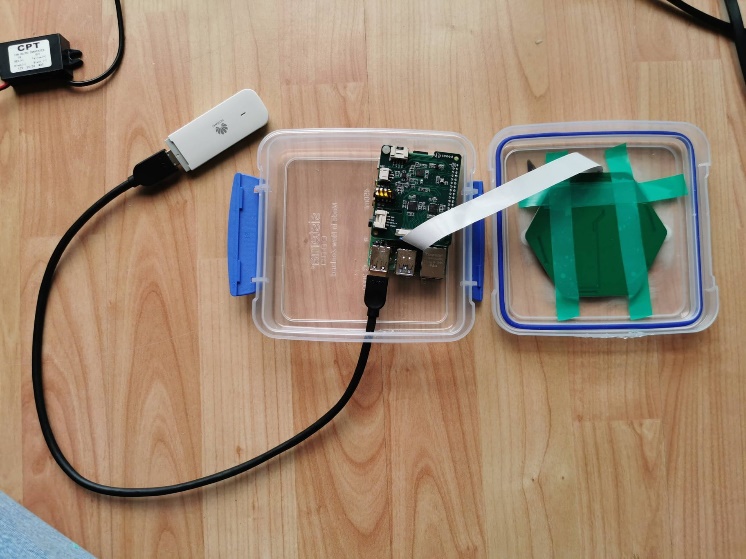 | 2. Put the raspberry Pi inside with the cables to the power source and the Huawei dongle coming out the bottom. Seal up the bottom with electric tape and clear sealant. |
| 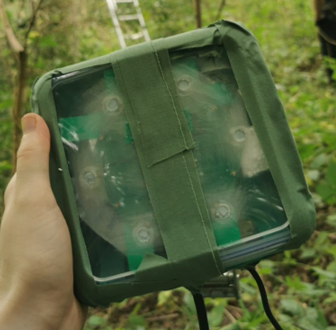 | 3. Put silica in the case and snap shut. Seal the edges with 2 layers of sniper tape. |
